# Small RNA modifications in Alzheimer’s disease

**DOI:** 10.1101/2020.04.30.071100

**Authors:** Xudong Zhang, Fatima Trebak, Lucas AC Souza, Junchao Shi, Tong Zhou, Patrick G. Kehoe, Qi Chen, Yumei Feng Earley

## Abstract

**Background:** While significant advances have been made in uncovering the aetiology of Alzheimer’s disease and related dementias at the genetic level, molecular events at the epigenetic level remain largely undefined. Emerging evidence indicates that small non-coding RNAs (sncRNAs) and their associated RNA modifications are important regulators of complex physiological and pathological processes, including aging, stress responses, and epigenetic inheritance. However, whether small RNAs and their modifications are altered in dementia is not known.

**Methods:** We performed LC-MS/MS–based, high-throughput assays of small RNA modifications in post-mortem samples of the prefrontal lobe cortices of Alzheimer’s disease (AD) and control individuals. We noted that some of the AD patients has co-occurring vascular cognitive impairment-related pathology (VaD).

**Findings:** We report altered small RNA modifications in AD samples compared with normal controls. The 15–25-nucleotide (nt) RNA fraction of these samples was enriched for microRNAs, whereas the 30–40-nt RNA fraction was enriched for tRNA-derived small RNAs (tsRNAs), rRNA-derived small RNAs (rsRNAs), and YRNA-derived small RNAs (ysRNAs). Interestingly, most of these altered RNA modifications were detected both in the AD and AD with co-occurring vascular dementia subjects. In addition, sequencing of small RNA in the 30–40-nt fraction from AD cortices revealed reductions in rsRNA-5S, tsRNA-Tyr, and tsRNA-Arg.

**Interpretation:** These data suggest that sncRNAs and their associated modifications are novel signals that may be linked to the pathogenesis and development of Alzheimer’s disease.

**Funding:** NIH grants (R01HL122770, R01HL091905, 1P20GM130459, R01HD092431, P50HD098593, GM103440), AHA grant (17IRG33370128), Sigmund Gestetner Foundation Fellowship to P Kehoe.

**Research in Context:** *Evidence before this study:* Alzheimer’s disease (AD) and vascular dementia (VaD) are marked by cognitive impairment and neuropathologies caused by significant neuronal death. Associated gene mutations are rare in subjects with dementia, and the aetiology of these diseases is still not completely understood. Recent emerging evidence suggests that epigenetic changes are risk factors for the development of dementia. However, studies assessing small RNA modifications—one of the features of epigenetics—in dementia are lacking.

*Added value of this study:* We used high-throughput liquid chromatography-tandem mass spectrometry and small RNA sequencing to profile small RNA modifications and the composition of small RNAs in post-mortem samples of the prefrontal cortex of AD and control subjects. We detected and quantified 16 types of RNA modifications and identified distinct small non-coding RNAs and modification signatures in AD subjects compared with controls.

*Implications of all the available evidence:* This study identified novel types and compositions of small RNA modifications in the prefrontal cortex of AD patients compared with control subjects in post-mortem samples. The cellular locations of these RNA modifications and whether they are drivers or outcomes of AD are still not known. However, results from the present study may open new possibilities for dissecting the dementia pathology.

## Introduction

Alzheimer’s disease (AD) and severe vascular cognitive impairment, also known as vascular dementia (VaD), are two of the most common types of dementia. They are characterized by cognitive impairment and neuropathology caused by neurotoxic forms of amyloid beta peptide, loss of synapses and neuronal function and, ultimately, significant neuronal loss. The aetiology of these forms of dementia is still not fully understood.^1^

Recent studies have highlighted epigenetic changes associated with aging or acquired through interactions with the environment as important risk factors for dementia, reflecting the fact that causative gene mutations are rare and are only present in a very small percentage of dementia patients. RNA modification is one of the most recently discovered epigenetic-mediated mechanisms for regulating physiological and pathological processes. Small RNA modifications, particularly those involving microRNAs (miRNAs)^2,3^ and transfer RNA-derived small RNAs (tsRNAs), have attracted considerable recent attention for their potential clinical relevance and possible use as diagnostic markers or therapeutic targets of diseases.^4–6^ To date, evidence for changes in small RNA modifications in dementia is lacking. Thus, to gain insights into possible new mechanisms and identify potential biomarkers for dementia, we undertook an exploratory study to profile small RNA expression and modification status in dementia.

In this report, we profiled small RNA modifications in post-mortem samples of human brain cortical tissue from patients with clinically and pathologically diagnosed AD and control individuals. To simultaneously detect and quantify 16 types of RNA modifications^7,8^ in different small RNA fractions [15–25 nucleotides (nt), 30–40 nt] from AD patients and control individuals, we used high-throughput liquid chromatography-tandem mass spectrometry (LC-MS/MS). We also profiled the composition of small RNAs in AD and control subjects using small RNA sequencing. These exploratory examinations revealed distinct sncRNA expression and RNA modification signatures between AD and controls, suggesting that these small RNAs are novel factors associated with the pathogenesis and/or progression of AD.

## Materials and methods

### Human subjects for RNA modification assays

A total of 34 post-mortem brain prefrontal lobe cortex samples from AD patients (n = 27) and control human subjects (n = 7) were collected by post-mortem autopsy, snap-frozen, and stored in liquid nitrogen. The time between biospecimen acquisition and analysis ranged from 5 to 30 years, according to the Brain Bank database. Biospecimens were shipped on dry ice. Information from clinical and pathology reports was obtained from the brain bank database, developed as described.^9^ On the basis of their clinical diagnosis, subjects were first categorized into two groups: no significant abnormalities (control) and AD. We realized that some AD patients had co-occurring vascular dementia. This type of patient usually has a more rapid course of disease clinically. We thus further divided the AD patients into AD without VaD (AD-w/o-VaD, n = 18) and AD with VaD (AD-w-VaD, n = 9) for the purpose of analysis. Subjects in the AD-w-VaD group were clinically diagnosed with clear AD tau pathology and Aβ plaques in post-mortem analysis by the UK Brain Bank, and a neuropathological report of AD. Subjects were selected according to their confirmed clinical and pathological diagnosis as AD or normal, and were obtained from the Human Tissue Authority-licensed South West Dementia Brain Bank http://www.bristol.ac.uk/translational-health-sciences/research/neurosciences/research/dementia/swdbb/, accessed on March 24^th^, 2020). detailed information of these subject is also summarized in Supplemental Table S1.

University of Bristol, with tissue bank ethics approval from the South West–Central Bristol Research Ethics Committee. Clinical and pathological data included patient history, diagnosis and medications; pathological reports were available for retrospective analysis. Patients’ personal information has been anonymized. Data and tissues for this study were collected between 1989 and 2015. The Research Integrity Offices at the University of Nevada, Reno, and University of Bristol have determined that this project complies with human research protection oversight by the Institutional Review Board.

### Isolation of total RNA from the human prefrontal cortex

Total RNA was isolated from 100 mg of human prefrontal tissues using the TRIzol reagent (Thermo Fisher Scientific, Inc. Waltham, MA, USA), as described by the manufacturer. RNA quantity was evaluated using a NanoDrop 2000 microvolume spectrophotometer (Thermo Fisher). RNA samples were stored at −80°C until analysis. Researchers performing small RNA sequencing and RNA modifications assays on RNA samples were blinded to diagnosis and group-identifying information.

### Small RNA Sequencing

Ten RNA samples from prefrontal lobe cortexes of control subjects and AD patients were submitted to BGI (Cambridge, MA, USA) for sequencing of 15–40-nt small RNAs. The number of clean reads varied from 30M to 40M across all samples. The Q20 score was greater than 99·5% for all samples. Raw sequencing data for each sample were processed and analysed using our newly developed computational framework, *SPORTS1·0*^1^ (https://github.com/junchaoshi/sports1.0), which is designed to optimize the annotation and quantification of non-canonical small RNAs (e.g., tsRNAs) in small RNA sequencing data. Sequencing adapters were removed, after which reads with lengths outside of the defined range and those with nucleotides other than ATUCG were discarded. Clean reads were mapped against the precompiled human small RNA annotation database (https://github.com/junchaoshi/sports1.0/blob/master/precompiled_annotation_database.md). Small RNAs 15–25 and 30–40 nt in length were summarized separately.

### Small RNA purification and analysis of RNA modifications using LC-MS/MS

Standardized ribonucleoside preparations, small RNA purification, and LC-MS/MS-based analysis of RNA modifications in RNA samples were performed as previously described.^8^ Purified small RNAs (100–200 ng) from human brain samples were digested as input and loaded onto a ThermoFisher Vantage Quadrupole mass spectrometer connected to a Thermo Ultimate 3000 UHPLC system equipped with an electrospray ionization source. The MS system was operated in positive ion mode using a multiple reaction monitoring (MRM) approach. LC-MS/MS raw data were acquired using Xcalibur Workstation software and were processed using Xcalibur QuanBrowser for quantification of modified ribonucleoside concentrations. We totally performed 34 samples. However, during the analysis of the mass spectrometry data, we exclude data points when the signal peak is not unique (e.g. two peaks, elevated baseline), or drifting off the detection time predetermined by each standardized ribonucleoside, which resulted in inaccurate reading (or loss of reading) of the examined signal. The percentage of each modified ribonucleoside was normalized to the total amount of quantified ribonucleosides containing the same nucleobase, an approach that decreases/eliminates errors caused by sample loading variation. For example, the percentage of m^5^C = mole concentration (m^5^C)/mole concentration (m^5^C + Cm + C + ac^4^C). Fold-changes in RNA modifications between different groups were calculated based on the percentage of modified ribonuclosides. Brain RNAs 15–25 and 30–40 nucleotides (nt) in length were examined.

### Statistical analysis

Data are presented as means ± SEM and were plotted using Prism8 software (GraphPad, La Jolla, CA, USA). Differential expression analyses were performed using the *edgeR* tool (pmid: 19910308), controlling for age and sex. RNAs with a false discovery rate < 10% were deemed to be differentially expressed. RNA modification levels among groups were compared using a linear model that controls for age and sex. Student *t*-test or Ordinary One-way ANOVA with post hoc Bonferroni correction was used as appropriate for comparisons among groups. A *P*-value < 0·05 was considered statistically significant.

## Results

### Participant characteristics

As summarized in Tables 1 and 2, patients and control individuals were similar in terms of age. Almost all patients were White/European except for one patient whose race was unknown. There were slightly more males (58·8%) than females (41·2%). As expected, brain weight was significantly lower in AD (*P* = 0·0099) compared with the controls (Table 1). This phenomenon persisted when we subdivided the AD into (AD-w/o-VaD) (*P* = 0·018) when compared with controls; however, there was no significant difference between AD-w-VaD and controls for the brain weight (Table 2). No significant differences were observed for the brain pH and post-mortem delay time among groups.

### Altered small RNA modification profiles in the cerebral cortex of AD patients

To systemically analyse small RNA expression and modification profiles, we collected the 15–40-nt RNA fraction from the brain frontal lobe cortex for small RNA sequencing as we previously described^7,8^, and then performed bioinformatic analyses using our recently developed software, *SPORTS1.0*.^10^. In addition, 15–25-nt and 30–40-nt RNA fractions were collected for high-throughput examination of RNA modification by LC-MS/MS (Supplemental Figure S1).^7,11^ These analyses resulted in the identification and quantification of 16 types of RNA modifications.

In the 15–25-nt RNA fraction from the cortex of AD brains (Figure 1A), we found an increase in 2’-O-methylcytidine (Cm), 7-methylguanosine (m^7^G), 2’-O-methylguanosine (Gm), and significant reductions in *N*^2^, *N*^2^, 7-trimethylguanosine (m^2,2,7^G) and *N*^2^, *N*^2^-dimethylguanosine (m^2,2^G), compared with controls (Figure 1B and 1C). Other RNA modifications that were not significantly different among groups were also detected and are shown in Supplemental Figure S2. Since AD Patients with co-occurring several vascular cognitive impairment-related pathologies (e.g. VaD) tend to have a more rapid course of disease clinically.^12^ We further test whether these patients have a distinct difference at RNA modifications. To this end, we divided AD patients into two groups, the AD-w/o-VaD and AD-w-VaD and performed further analysis. We found that both AD-w/o-VaD group and AD-w-VaD group showed similar trends in RNA modification changes when compare to the normal subjects, but some RNA modifications cannot reach statistical significance when the two groups are divided (Figure 2), possibly due to decreased sample sized in both groups.

**Figure 1.**
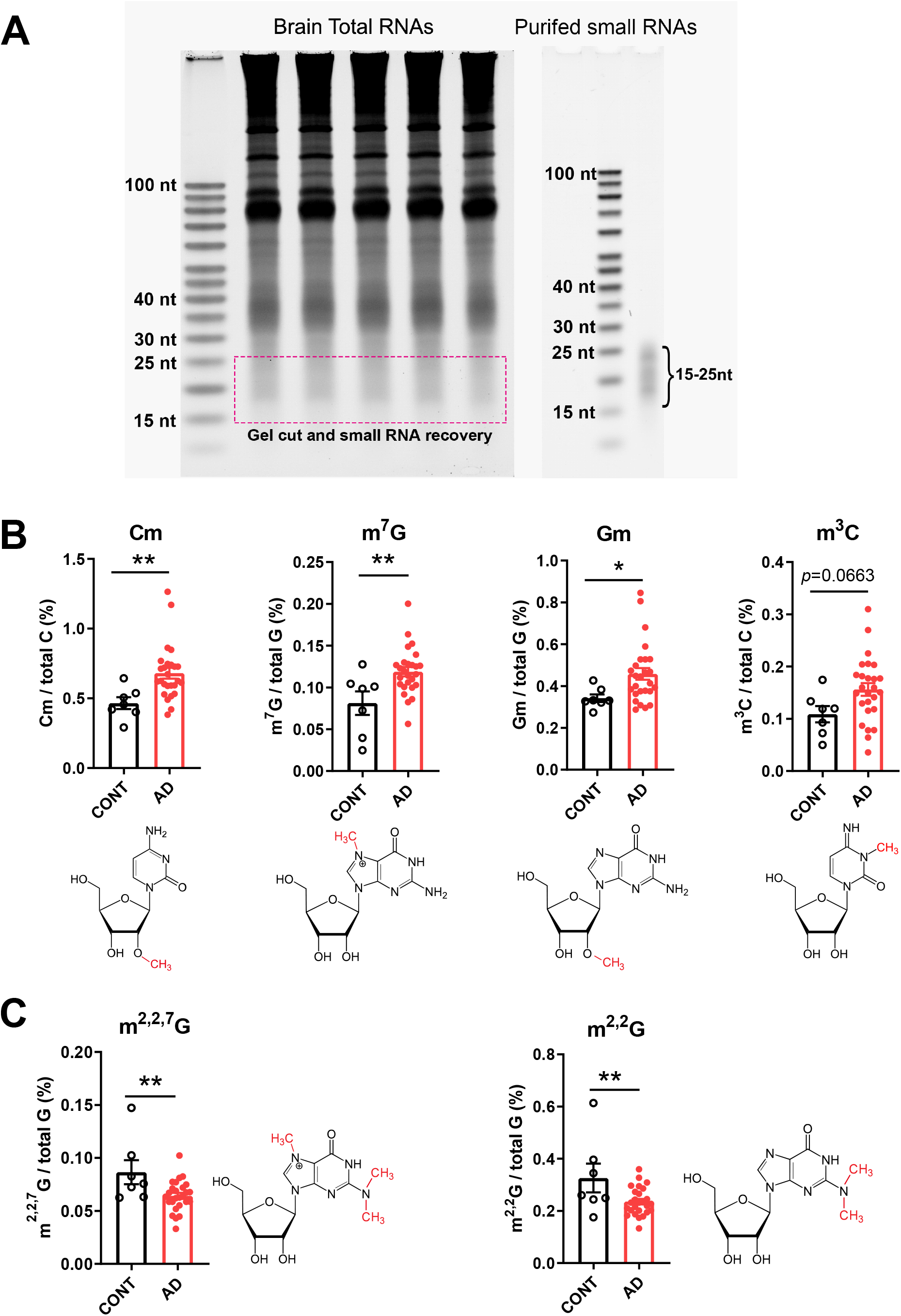
Small RNA modifications in the 15–25-nt fraction from samples of the prefrontal lobe cortex of dementia patients. (**A**) Representative image showing purification of 15–25-nt small RNAs for the analysis of RNA modifications. (**B**) Increases in 2’-O-methylcytidine (Cm) 7-methylguanosine (m^7^G), 2’-O-methylguanosine (Gm), and 3-Methylcytidine(m^3^C) modifications. (**C**) Reductions in *N*^2^,*N*^2^,7-trimethylguanosine (m^2,2,7^G) and *N*^2^,*N*^2^-dimethylguanosine (m^2,2^G) modifications in the cortex of AD compared with CONT subjects (each dot in the figure represents value from one independent sample); \**P* < 0·05;\*\**P* <0·01 vs. CONT by Student *t*-test.

**Figure 2.**
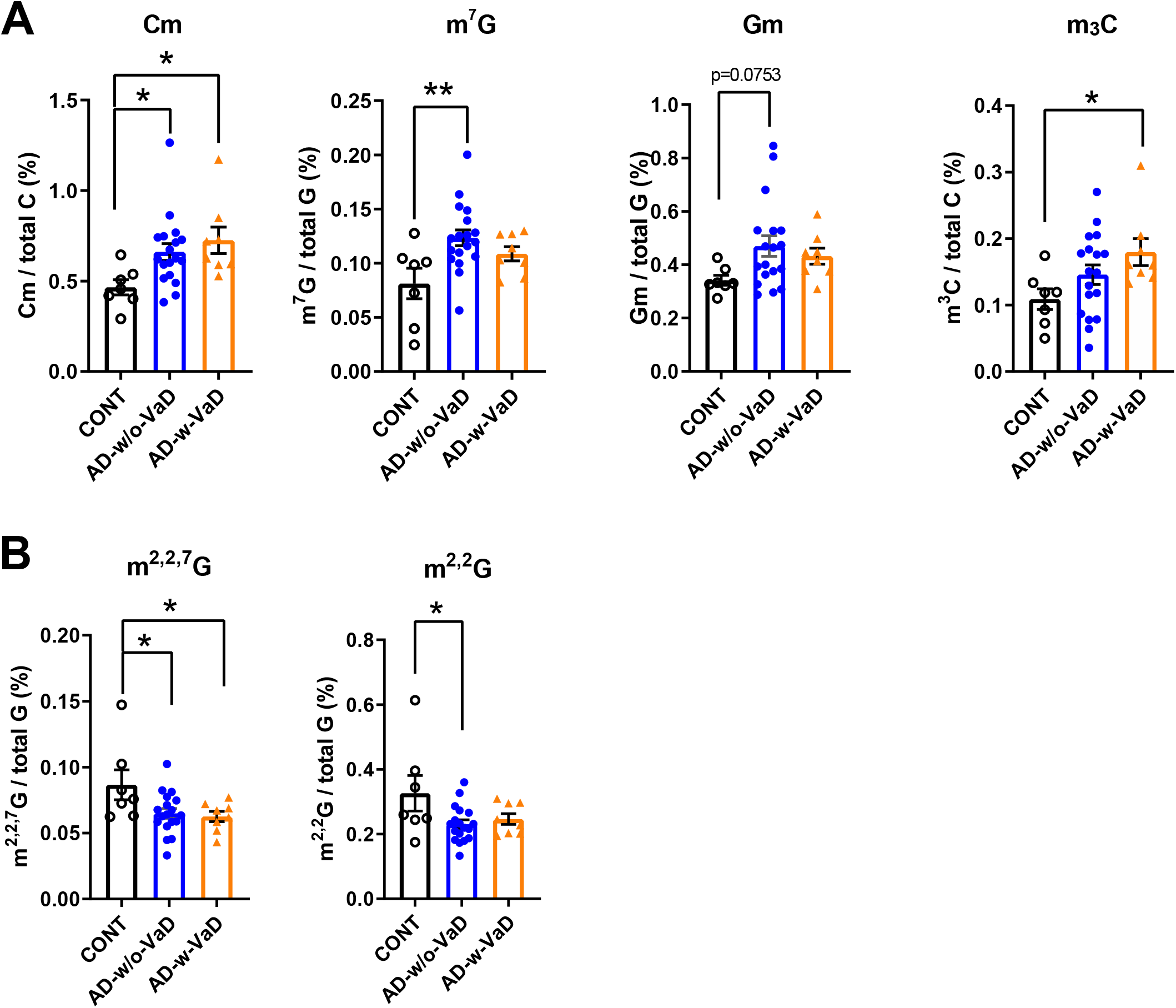
Small RNA modifications in the 15–25-nt fraction from samples of the prefrontal lobe cortex of dementia patients with and without co-occurring vascular dementia patients. (**A**) Increases in 2’-O-methylcytidine (Cm) 7-methylguanosine (m^7^G), 2’-O-methylguanosine (Gm), and 3-Methylcytidine(m^3^C) modifications. (**B**) Reductions in *N*^2^,*N*^2^,7-trimethylguanosine (m^2,2,7^G) and *N*^2^,*N*^2^-dimethylguanosine (m^2,2^G) modifications in the cortex of AD-w/o-VaD, AD-w-VaD, and CONT subjects (each dot in the figure represents value from one independent sample); \**P* < 0·05;\*\**P* <0·01 vs. CONT by Ordinary One-way ANOVA with post hoc Bonferroni correction.

In the 30–40-nt small RNA fraction from the cortex of AD brains (Figure 3A), we found significantly higher levels of Cm, 2’-O-methyluridine (Um) and 7-methylguanosine (m^7^G) modifications, and reductions in 1-methylguanosine (m^1^G), m^2,2,7^G and pseudouridine (psi or Ψ) modifications compared with controls (Figure 3B and 3C). Other RNA modifications detected in this 30–40-nt fraction that did not exhibit significant differences among groups are shown in Supplemental Figure S3. When further divided AD into with or without VaD (Figure 4), we found that AD-w/o-VaD maintained significant changes in most of these RNA modifications. One the other hand, the AD-w-VaD group maintained the same trend but did not reach significant difference in most of the RNA modifications compared with the controls possibly due to the small sample size; however, there was no difference between the AD-w-VaD and the AD-w/o-VaD suggesting the similar expression profile.

**Figure 3.**
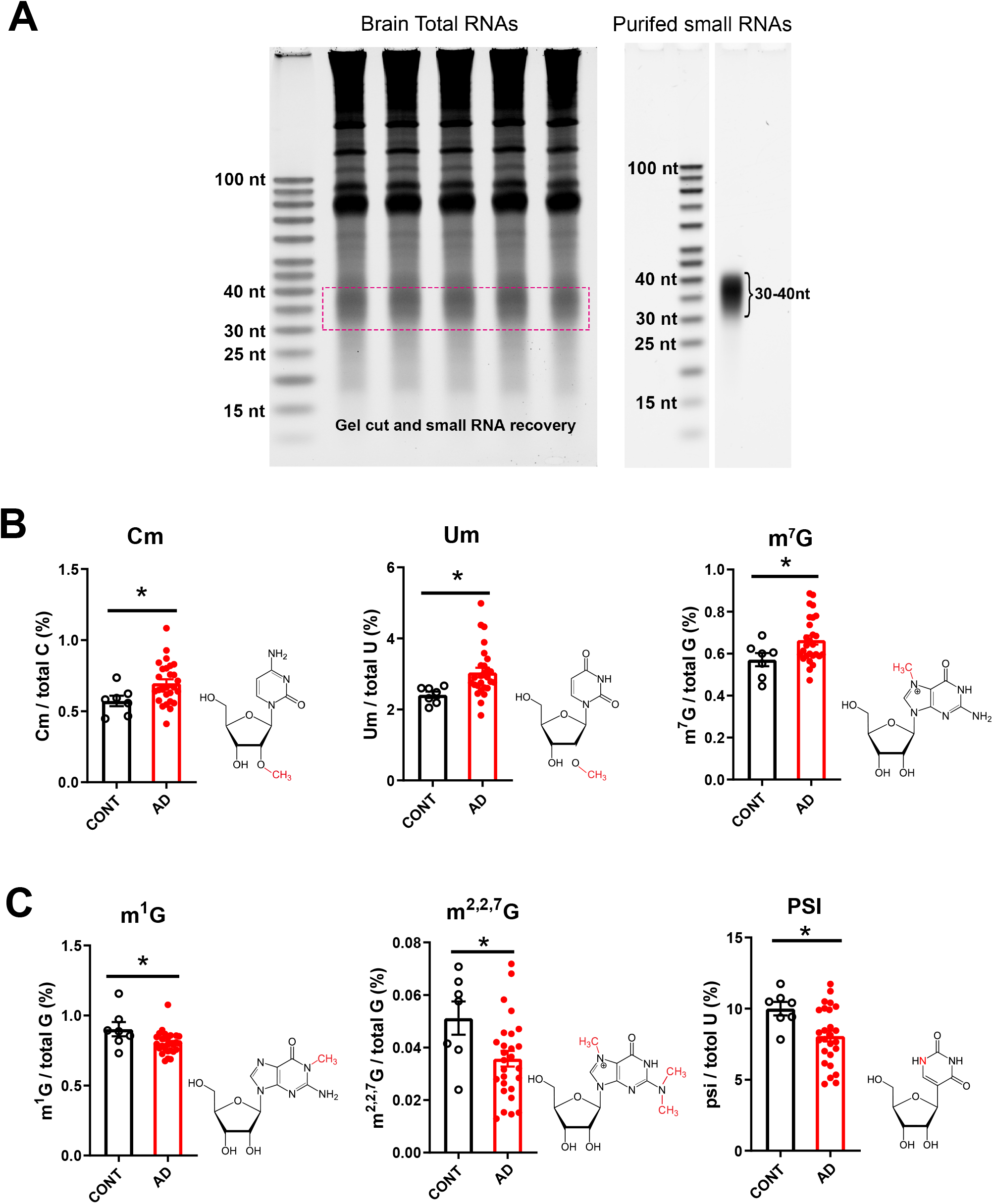
Small RNA modifications in the 30–40-nt fraction from samples of the prefrontal lobe cortex of dementia patients. (**A**) Representative image showing purification of 30–40-nt small RNAs for the analysis of RNA modifications. (**B**) Increases in 2’-O-methylcytidine (Cm), 2’-O-methyluridine (Um) and 7-methylguanosine (m^7^G) modifications, and (**C**) reductions in 1-methylguanosine (m^1^G), *N*^2^,*N*^2^-dimethylguanosine (m^2,2^G) and pseudouridine (psi or ψ) modifications in the cortex of AD compared with CONT subjects (each dot in the figure represents value from one independent sample); \**P* < 0·05 vs. CONT by Student *t*-test.

**Figure 4.**
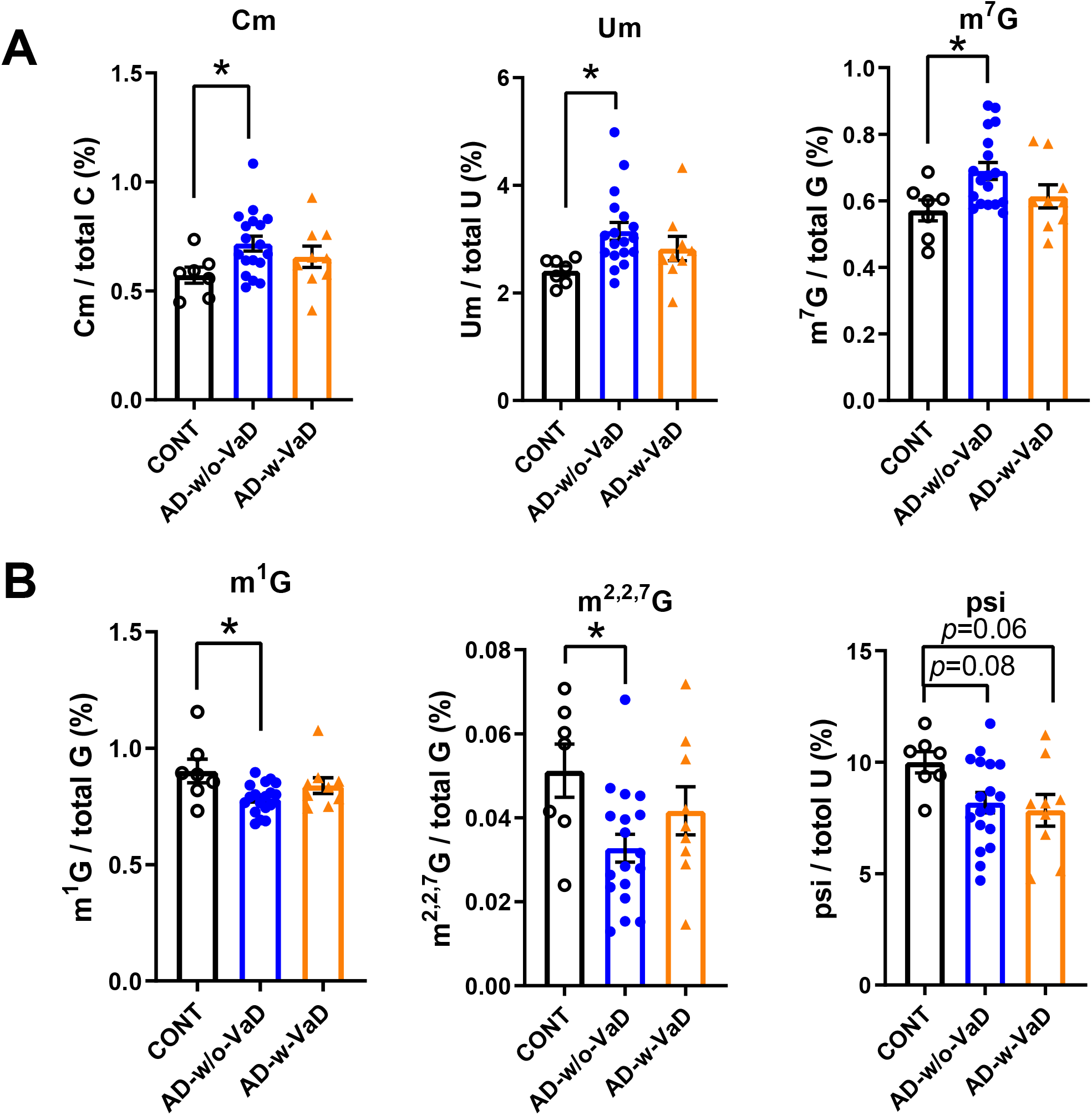
Small RNA modifications in the 30–40-nt fraction from samples of the prefrontal lobe cortex of dementia patients with and without co-occurring vascular dementia patients. Changes in (**A**) 2’-O-methylcytidine (Cm), 2’-O-methyluridine (Um) and 7-methylguanosine (m^7^G) modifications, and (**B**) 1-methylguanosine (m^1^G), *N*^2^,*N*^2^-dimethylguanosine (m^2,2^G) and pseudouridine (psi or ψ) modifications in the cortex of AD-w/o-VaD, AD-w-VaD, and CONT subjects (each dot in the figure represents value from one independent sample); \**P* < 0·05 vs. CONT by Ordinary One-way ANOVA with post hoc Bonferroni correction.

### Altered 30–40-nt small RNA expression profiles in the cerebral cortex of AD patients

To systemically analyse small RNA expression profiles, we collected the 15–40-nt fraction from the brain frontal lobe cortex of AD patients and controls for small RNA sequencing, as we previously described^7,8^, and then performed bioinformatic analyses using our recently developed software, *SPORTS1·0*^10^. Our RNA-seq analysis showed that the 15–25-nt RNA population was predominantly miRNAs (Figure 5A) in both the AD and control samples; we did not find significant differences in specific small RNA sequences in this fraction between AD and control samples (data not shown). Whereas the expression profile of the small RNA population in the 30–40-nt fraction showed more dynamic changes. (Figure 5B-E). Our RNA-seq data revealed three major subtypes in the 30–40-nt fraction, namely tRNA-derived small RNAs (tsRNA), rRNA-derived small RNAs (rsRNAs) and Y RNA-derived small RNAs (ysRNAs); other un-annotated RNAs were also found (Figure 5B). Although there were trends toward an increase in tsRNAs and a reduction in ysRNAs in AD patients, these differences did not reach statistical significance. Interestingly, in the 30–40-nt fraction, we found a significant reduction in rsRNA-5S, tsRNA-Tyr and tsRNA-Arg fragments in AD patients compared with controls (Figure 5C–E).

**Figure 5.**
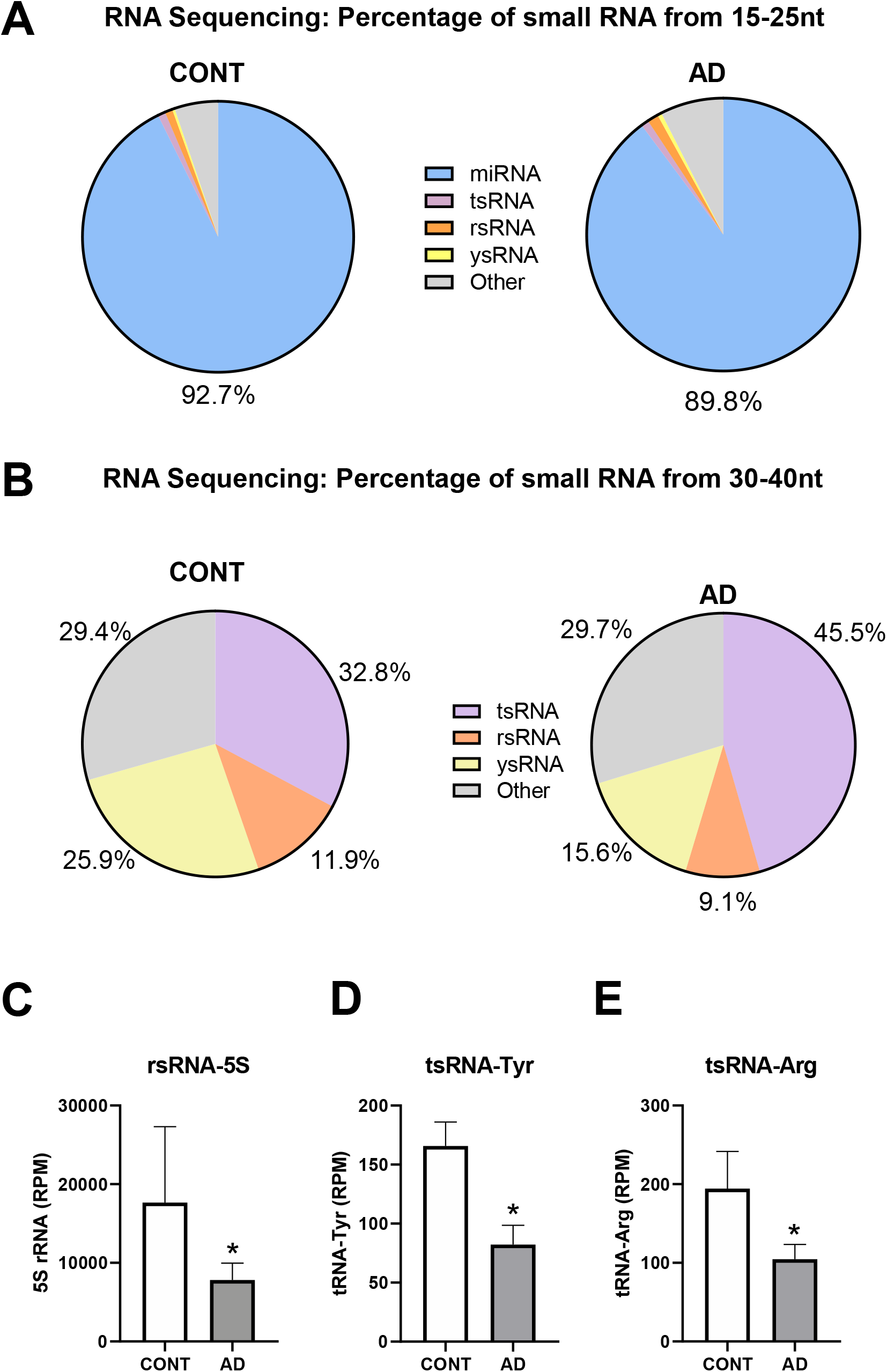
Reductions in rsRNA-5S, tsRNA-Tyr, and tsRNA-Arg in prefrontal lobe cortex samples of AD patients, identified by small RNA sequencing. (**A**) Composition of the 15–25 nt fraction of small RNAs. (**B**) Composition of the 34–40 nt fraction of small RNAs. (**C**) rsRNA-5S, (**D**) tsRNA-Tyr, (**E**) tsRNA-Arg in the cortex of AD patients compared with CONT subjects (n = 4 for CONT, n = 6 for AD in panel A-E); \**P* < 0·05 vs. CONT by Student *t*-test.

## Discussion

Emerging evidence shows that small RNAs, including miRNAs, tsRNAs and piwi-interacting RNAs, harbour a diversity of RNA modifications.^13^ The physiological and pathological importance of these small, non-coding RNA modifications has only recently begun to emerge, as highlighted by studies revealing their active involvement in stress responses, metabolism, immunity, and epigenetic inheritance of environmentally acquired traits.^4,14,15^ RNA modifications have recently been detected in the nervous system, where they are involved in the regulation of cortical differentiation, behaviour, and brain functions.^16,17^ However, to our knowledge, there have been no reports on the status of small RNA modifications in AD or VaD to date.

The current study showed intriguing differences in small RNA-modification signatures between AD patients and control subjects, suggesting an association and possible contribution of such modifications to the pathogenesis and/or progression of AD. Notably, most of the altered RNA modifications were observed in the AD with and without VaD, providing molecular evidence for potential distinct signatures for AD patients. In the last three decades, dementia research has demonstrated significant overlaps between AD and VaD in terms of clinical symptoms, risk factors, and post-mortem brain autopsy findings.^18–20^ Therapeutically, drugs that enhance cholinergic activity are as effective in patients suffering from VaD as they are in AD patients.^21^ More recently, drugs that have traditionally been used for cardiovascular diseases, especially renin-angiotensin system blockers, have shown benefits not only in VaD patients, but also in AD patients.^22–24^ However, the molecular pathogenesis of AD with co-occurring sever vascular cognitive impairment remains incompletely understood. It is however worth highlighting that patients with AD and co-occurring severe Vascular Cognitive Impairment-related pathology (e.g. VaD) tend to have a more rapid course of disease clinically because of the co-occurrence of heightened cerebrovascular disease and hypoxia and related white matter and neuronal damage.^12,25^ It was therefore prudent to consider these as a separate group to explore whether RNA modifications might be more prevalent and thus potentially associated with the disease. We were not surprised to find many common signatures among RNA modifications in AD with or without VaD patients, particularly in the 15–25-nt small RNA modifications that might contribute to shared mechanisms of dementia pathology. Yet, the lack of apparent differences between the groups does suggest, notwithstanding the small sample sizes, that the RNA modifications may relate more to AD-related processes than neuropathology caused by VaD. Further studies, involving post-mortem analysis of other cerebrovascular-related diseases and in the absence of AD or with minimal pathology, and with larger sample size, would be useful studies to explore this further.

The 15–25-nt RNAs were predominantly miRNAs, whereas 30–40-nt RNAs were predominantly tsRNAs, rsRNAs and ysRNAs, as demonstrated by small RNA sequencing. We found changes in RNA-modification profiles in the AD brain cortex in both fractions of small RNAs. While the involvement of miRNA in the pathogenesis of AD has been explored^2,3,26^, the identifications of rsRNAs and ysRNAs in a pathophysiological context has only begun to emerge^8,27,28^ and has not been studied in AD. The dynamic expression of these non-canonical small RNAs suggests the possibility of unidentified biological functions that warrant further investigation^29,30^, particularly in relation to several risk factor genes that have been identified for AD in the last few decades.^31,32^ This situation is similar to that for tsRNAs, which have shown recently expanding functions.^6,33,34^ The novel RNA modifications that showed dynamic changes in AD patients included Cm, Um, m^7^G, m^1^G, m^3^C, m^2,2^G, m^2,2,7^G, and psi. The functions and identities of these small RNA modifications in AD are not yet understood, but our data suggest that these RNA modifications in the brain cortex of dementia patients are potentially important, either as a consequence or an actual cause of pathogenesis. For example, it has been reported that pH, which may affect the efficiency of related RNA-modifying enzymes and thus contribute to the RNA modification profile, is lower in the brains of AD patients compared with normal aging controls^35,36^. Although we observed a trend toward lower pH values in post-mortem samples from AD w/o VaD brains, this difference did not reach statistical significance.

tRNA-derived small RNAs are reported to contribute to multiple pathological processes, including cancer, viral infection, and age-related neurodegenerative diseases.^4^ In the senescence-accelerated mouse prone 8 (SAMP8) model, which mimics age-related neuro-disorders such as AD and Parkinson disease, tsRNAs, including tsRNA-Tyr and tsRNA-Arg, in brain tissue are dysregulated compared with normal brain tissue. The targets of these dysregulated tsRNAs are enriched in neurodevelopment pathways such as synapse formation.^37^ In addition, knockout of the RNA kinase, CLP1, in mice leads to progressive loss of motor neuron function in association with the accumulation of tsRNAs derived from Try-tRNA and Arg-tRNA.^38,39^ These tsRNAs sensitize motor neurons to oxidative stress-induced cell death, suggesting that tsRNAs are involved in normal motor neuron functions and responses to oxidative stress.^38^ A similar phenomenon was reported in a human neurological disease cohort, and it was shown that transfection of small RNAs derived from 5’ Tyr-tRNA can protect CLP-mutant cells from oxidative stress-induced cell death.^39^ In our study, the decrease in 5’tsRNA-Tyr in AD subjects may confer greater vulnerability to oxidative stress on neurons. Interestingly another report showed that the expression of 5S rRNA and levels of oxidized 5S rRNA are dynamically modulated in AD’s subjects^40^, findings that may be related to the biogenesis of rsRNAs and could explain alterations of rsRNA-5S in the AD group in the current study.

Whether the changes in small RNAs and RNA modifications observed here represent drivers or outcomes of AD remains unknown. In our study, the sample number from the control and patient groups is relatively small, which prevent deeper analyses to consider other factors such as ages, NFT pathology stage, and PMI. One limitation of the current study is the prolonged PMI in some of the brain samples as we have no access to adequate number of samples with shorter PMI (e.g. < 24 hours PMI). Another potential limitation of our study is the prolonged storage time in some of the tissues, as long-term storage might affect the RNA modification results. To examine this possibility. We performed principle components analysis (PCA), which converts the observations of possibly correlated variables (RNA modifications) into a set of values of linearly uncorrelated variables. This analysis of the 27 AD samples ranging from 1985 to 2015 does not support this possibility that storage time affects RNA modification as showed in Supplemental Figure S4. This result support the conclusion that the detected changes in RNA modifications indeed represent intrinsic difference in the AD samples and the storage time is not a major contributing factor on the result.

In summary, the current pioneer study based on limited sample numbers have revealed that RNA modifications in the sncRNAs are associated with AD, although the precise meaning of each of these changes in terms of their specific functions awaits discovery. Future efforts to pinpoint the locations of these modifications in each sncRNA population and identify enzymes involved in their regulation would be invaluable^41,42^ and will open new avenues for dissecting the nature of dementia pathology.

## Data availability

Small RNA-seq data that support the findings of this study have been deposited in the Gene Expression Omnibus (GEO) under accession code GSE153284. All other data supporting the findings of this study are available from the corresponding author on reasonable request.

## Acknowledgements

We would like to thank Dr. David Quilici and Rebekah Woolsey from the Mick Hitchcock, Ph.D. Nevada Proteomics Center for technical support with HPLC/MS instruments. We would especially like to thank Dr. Laura Palmer for tissue processing and facilitating access to clinical data from the South West Dementia Brain Bank at the University of Bristol, which is supported by grants from BRACE, ABBUK.

