## Supplemental Figure for "Small RNA modifications in Alzheimer’s disease" (from another version of this preprint: version 1, 2020-05-01)

### Title: Evidence for unique small RNA modifications in Alzheimer's disease

#### (Data Supplement)

### Figure S1.

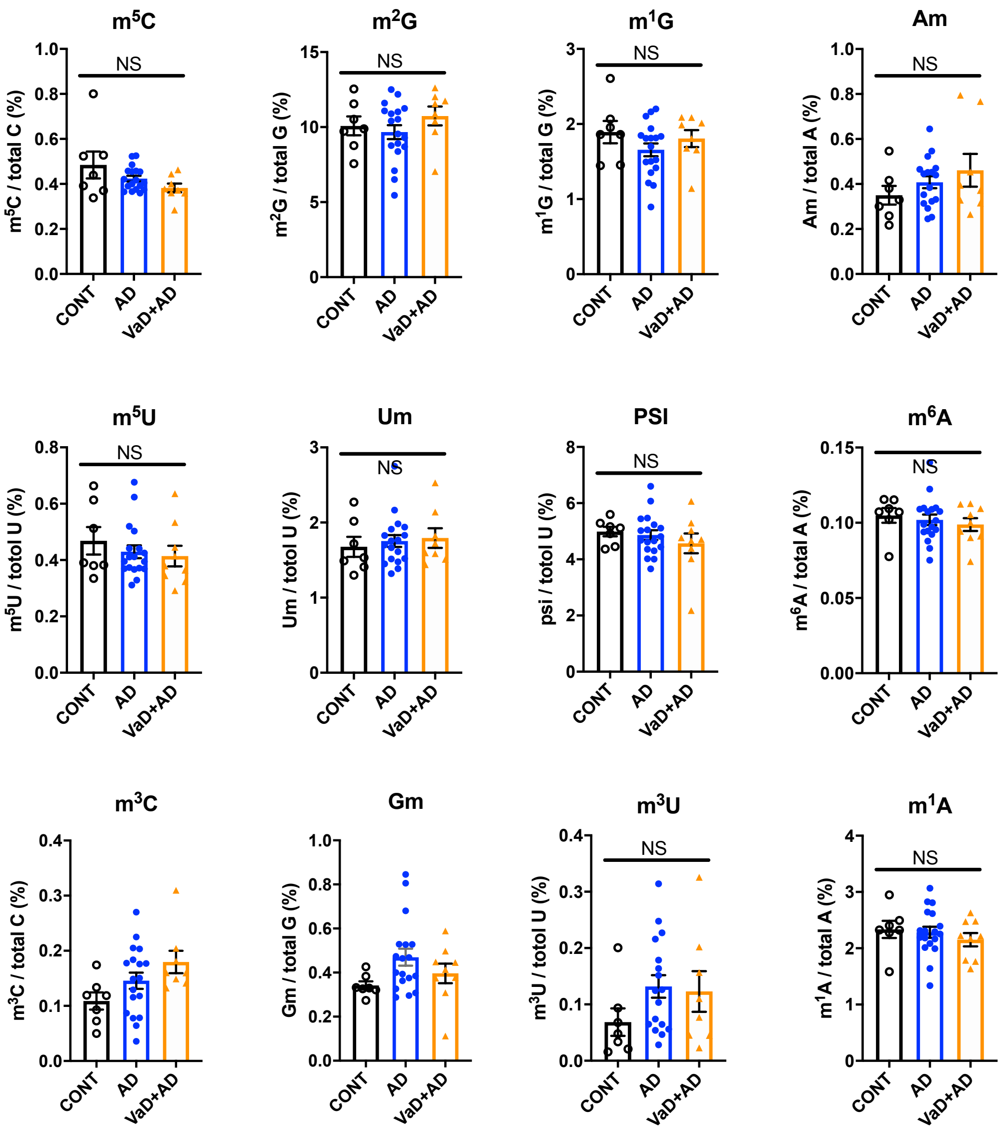


**Figure S1.** miRNA modifications in the prefrontal lobe cortex of AD and VaD+AD patients. There were no changes in 15–25-nt small RNA modifications in the cortex of AD and VaD+AD compared with CONT subjects (n = 7 for CONT, n = 18 for AD, and n = 9 for VaD+AD; * $P < 0{\cdot}05$ vs. CONT).

### Figure S2.

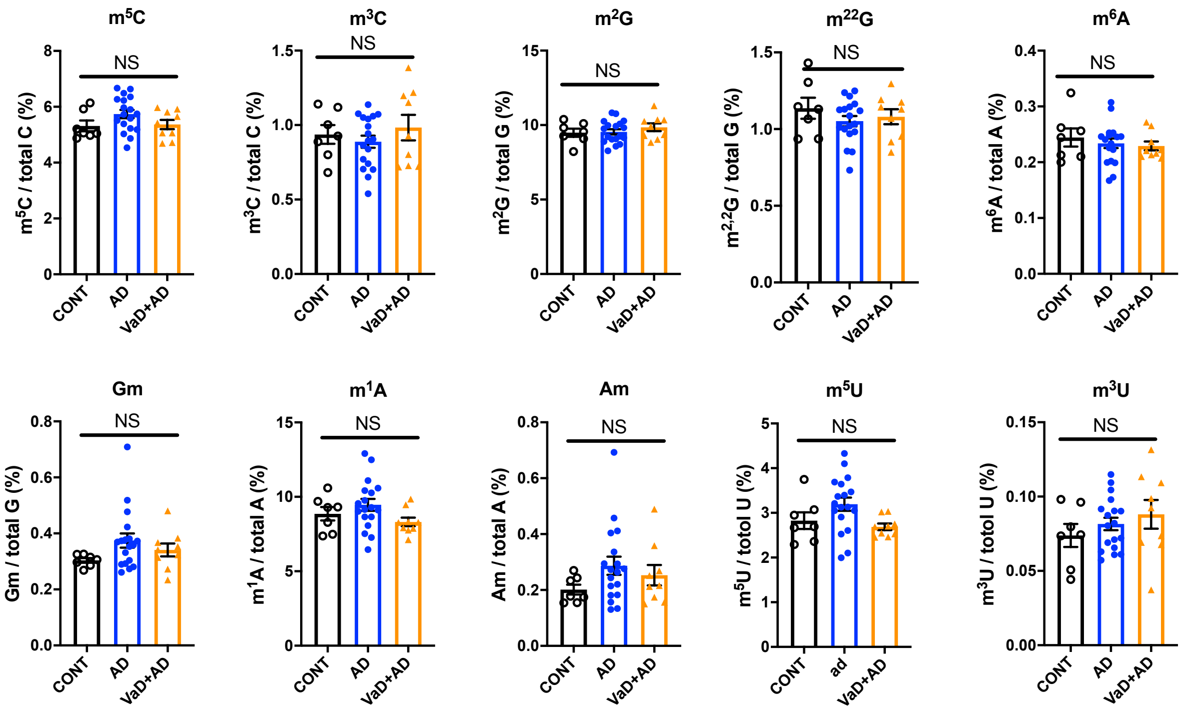


**Figure S2.** Small RNA (30–40 nt) modifications in the prefrontal lobe cortex of AD and VaD+AD patients. There were no changes in 30–40-nt small RNA modifications in the cortex of AD and VaD+AD compared with CONT subjects (n = 7 for CONT, n = 18 for AD, and n = 9 for VaD+AD; * $P < 0{\cdot}05$ vs. CONT).
